# Selective co-activation of α7- and α4β2-nicotinic acetylcholine receptors reverses beta-amyloid-induced synaptic dysfunction

**DOI:** 10.1101/2020.11.05.370080

**Authors:** Jessica P. Roberts, Sarah A. Stokoe, Matheus F. Sathler, Robert A. Nichols, Seonil Kim

## Abstract

Beta-amyloid (Aβ) has been recognized as an early trigger in the pathogenesis of Alzheimer’s disease (AD) leading to synaptic and cognitive impairments. Aβ can alter neuronal signaling through interactions with nicotinic acetylcholine receptors (nAChRs), contributing to synaptic dysfunction in AD. The three major nAChR subtypes in the hippocampus are composed of α7-, α4β2-, and α3β4-nAChRs. Aβ selectively affects α7- and α4β2-nAChRs, but not α3β4-nAChRs in hippocampal neurons, resulting in neuronal hyperexcitation. However, how nAChR subtype selectivity for Aβ affects synaptic function in AD is not completely understood. Here, we showed that Aβ associated with α7- and α4-containing nAChRs but not α3-containing receptors. Computational modeling suggested two amino acids in α7-nAChRs, Arginine 208 and Glutamate 211, were important for the interaction between Aβ and α7-containing nAChRs. These residues were found to be conserved only in the α7 and α4 subunits. We therefore mutated these amino acids in α7-containing nAChRs to mimic the α3 subunit and found that mutant α7-containing receptors were unable to interact with Aβ, providing direct molecular evidence for how Aβ selectively interacted with α7- and α4-containing receptors, but not α3-containing nAChRs. Selective co-activation of α7- and α4β2-nAChRs was also sufficient to reverse Aβ-induced AMPA receptor (AMPAR) dysfunction, including Aβ-induced reduction of AMPAR phosphorylation and surface expression in hippocampal neurons. Moreover, the Aβ-induced disruption of long-term potentiation was reversed by co-stimulation of α7- and α4β2-nAChRs. These findings support a novel mechanism for Aβ’s impact on synaptic function in AD, namely the differential regulation of nAChR subtypes.

## Introduction

Alzheimer’s disease (AD) is the predominant cause of dementia in the elderly, which is characterized by two histopathological hallmarks, beta-amyloid peptide (Aβ)-containing senile plaques and hyperphosphorylated tau-based neurofibrillary tangles (1). One of the early cognitive symptoms of AD is hippocampus-dependent memory impairments (2). Although neurodegeneration in AD is associated with multiple cellular abnormalities including tauopathies, mitochondrial dysfunction and oxidative stress, neuroinflammation, and gliosis (3,4), many studies have provided evidence that oligomeric Aβ triggers synaptic dysfunction and loss of hippocampus-dependent memory in AD (4–6). In particular, accumulation of Aβ in the prodromic stage of AD is strongly associated with Aβ’s contribution to the synaptic dysfunction (6,7). Although deficits in many neurotransmitter systems, including γ-aminobutyric acid (GABA) and serotonin, are associated with the progression of AD, the early symptoms appear to correlate strongly with dysfunction of cholinergic and glutamatergic synapses (6). However, the precise mechanisms of Aβ-induced deficits in these synapses remain to be determined.

The cholinergic system has been postulated to be a primary target in AD (8). A loss of cholinergic function is strongly associated with the onset of memory deficits in AD (9). Specifically, the loss of basal forebrain cholinergic neurons and altered nicotinic acetylcholine receptor (nAChR) expression in multiple regions of the brain, including in the hippocampus, are prominent pathological hallmarks in AD (10–12). In contrast, the expression of most muscarinic acetylcholine receptors (mAChR) subtypes is relatively unaltered in AD (13,14). The nAChR-mediated cholinergic modulation of hippocampal synaptic plasticity, such as long-term potentiation (LTP) and long-term depression (LTD), plays a critical role in learning and memory (15). Importantly, cholinergic synapses in the hippocampus are impaired by Aβ in the early stage of AD (16). Indeed, Aβ can alter neuronal signaling through interactions with nAChRs, ultimately contributing to synaptic dysfunction in AD (reviewed in (17). There are diverse lines of evidence that molecular interactions between Aβ and nAChRs affect receptor function in the early stages of AD (18–20). Nonetheless, contradictory results have been reported describing the effects of Aβ on nAChR physiology. For example, Aβ has been reported to bind to these receptors and produce functional receptor activation or inhibitory effects, depending on Aβ concentration, type of preparation (i.e., monomers, soluble oligomers or fibrils) and incubation times (21–23). Therefore, there is a need to determine how Aβ specifically affects nAChRs and contributes to AD pathogenesis.

While nearly 30 subtypes of neuronal nAChRs have been reported, the three major nAChR subtypes in the hippocampus are composed of α7, α4β2, and α3β4 subunits (24–26). Importantly, most of the current U.S. Food and Drug Administration (FDA)-approved drugs for AD inhibit the general breakdown of acetylcholine (acetylcholinesterase inhibitors), which potentially stimulates all types of nAChRs. Thus, it is not surprising that these receptors modulators are only moderately effective (12,27,28). In addition, the observation that Aβ accumulates in brain regions enriched for α4β2- and α7-nAChRs may provide an important clue for the selective vulnerability of the hippocampus to Aβ toxicity given the high-affinity interaction between Aβ and these nAChRs (29–32).

Our previous work using Ca^2+^ imaging in cultured hippocampal neurons has shown that Aβ selectively inhibits α7- and α4β2-nAChRs together, but not α3β4-nAChRs (32), indicating that distinct nAChR subtypes are differentially affected in AD. As nAChRs are more prominently expressed in inhibitory interneurons than excitatory cells in the hippocampus (33,34), nAChR-mediated cholinergic activity in the hippocampus may be biased toward altering the excitability of inhibitory interneurons. In line with this idea, our previous work demonstrates that Aβ induces neuronal hyperexcitation, an important characteristic in AD linked to network hyperexcitability and consequential dysfunction in brain rhythms (5), in cultured hippocampal excitatory neurons by predominantly reducing neuronal activity in inhibitory neurons via selective inhibition of α7- and α4β2-nAChRs, but not α3β4-nAChRs (32). Consistent with these findings, considerable evidence suggests that Aβ exerts subtype-specific inhibition of α7- and/or α4β2-nAChR function without affecting α3β4-nAChRs (21–23,32,35–39). The expression of α7 and α4 subtypes is also more significantly reduced in the cortex and hippocampus of AD patients compared to α3-type receptors (40,41). This suggests Aβ-induced disruption of selective nAChR function may induce synaptic and neuronal dysfunction in the hippocampus, leading to cognitive decline in AD. Therefore, strategies that selectively regulate nAChRs in the hippocampus can reverse the pathological Aβ effects on AD pathology, which may improve cognitive function. However, how nAChR subtype selectivity of Aβ affects synaptic function in AD is not completely understood.

Previous work using structure-function analysis has shown that the hydrophilic N-terminal domain of Aβ affects α7- and α4β2-nAChR function, elevating presynaptic Ca^2+^ levels in a model reconstituted rodent neuroblastoma cell line and isolated mouse nerve terminals (42). Furthermore, the activity of the Aβ N-terminus largely comes from a sequence surrounding a putative Histidine-based metal binding site, YEVHHQ (42). More importantly, this hexapeptide Aβ core sequence (Aβcore) is found to dock into the ligand binding site of nAChRs and reverse Aβ-induced neuronal apoptotic death, synaptic plasticity and fear memory deficits (43). In addition, mutations of Tyrosine to Serine and the two Histidine residues to Alanines in Aβcore (SEVAAQ) substantially reduce its neuroprotective effects, identifying these residues as critical to the neuroprotective actions of the Aβcore (43). These findings are consistent with earlier evidence showing that a different core Aβ fragment, Aβ_12-28_, that contains the critical residues of the Aβcore is sufficient to prevent Aβ from binding to α7-nAChRs and reverse Aβ-induced inhibition of α7-nAChRs (44). Finally, a recent study shows that the formation of Aβ-α4β2-nAChRs complex is based on the interaction of a part of Aβcore sequence (EVHH) with α4β2-containing receptors, and blocking this interaction prevents Aβ42-induced inhibition of α4β2-nAChRs (45). This thus suggests interrupting the association of Aβ with nAChRs may be neuroprotective against Aβ-induced neuronal dysfunction in AD, although the differential impact of Aβcore on the three major nAChRs in the hippocampus remains to be explored.

Here, we investigated the Aβ interaction with nAChRs in Aβ-induced Ca^2+^ hyperexcitation in cultured hippocampal neurons, assessing the impact of the neuroprotective, non-toxic N-terminal Aβ core peptide (Aβcore) to reverse the hyperexcitation. In addition, we assessed the selective interaction of Aβ with specific nAChRs, while identifying the amino acids, Arginine and Glutamate, within the loop C of the α7 and α4 subunits critical for these interactions. Moreover, we examined the impact of selective co-activation of α7- and α4β2-nAChRs on Aβ-induced synaptic dysfunction, including regulation of AMPA-type glutamate receptors. The findings have implications for regulation of nAChRs as therapeutic targets in the hippocampus for neuroprotection in AD.

## Results

### Interaction between Aβ and nAChRs in Aβ-induced Ca^2+^ hyperexcitation

Altering the interaction of Aβ with α7- and α4β2-nAChRs may be neuroprotective against Aβ-induced neuronal dysfunction in AD. We thus examined whether the interaction of Aβ_1-42_ (Aβ42) with nAChRs is important for neuronal hyperactivity by using the Aβcore peptide (42,43). As neuronal Ca^2+^ indicates neuronal activity (46), we measured Ca^2+^ activity in cultured 12-14 days *in vitro* (DIV) mouse hippocampal pyramidal neurons transfected with GCaMP6f (a genetically encoded Ca^2+^ indicator) as described previously (32,47). We treated neurons with soluble Aβ42 oligomers (oAβ42) and determined Ca^2+^ activity in hippocampal neurons immediately after treatment. We found active spontaneous Ca^2+^ transients in the control condition (250nM scrambled Aβ42; sAβ42) (**Fig. 1**). Consistent with the previous findings (32), total Ca^2+^ activity in 250nM oAβ42-treated cells was significantly higher than in sAβ42-treated controls (sAβ42, 1.00±0.65 ΔF/F_min_ and oAβ42, 1.48±0.95 ΔF/F_min_, *p* = 0.0009), confirming that soluble 250nM Aβ42 oligomers were sufficient to increase neuronal Ca^2+^ activity (**Fig. 1**). Notably, when 1μM of Aβcore was added in conjunction with 250nM oAβ42, Aβcore treatment was able to reverse oAβ42-induced Ca^2+^ hyperexcitation (oAβ42+Aβcore, 0.98±0.93 ΔF/F_min_, *p* = 0.04) (**Fig. 1**). However, the Aβcore had no effect on GCaMP6f activity in sAβ42-treated control neurons (sAβ42+Aβcore, 0.85±0.48 ΔF/F_min_) (**Fig. 1**), which is consistent with the previous finding that Aβcore treatment had no prolonged effect on Ca^2+^ levels in differentiated mouse neuroblastoma cells (43). Next, we added 1μM inactive Aβcore in oAβ42-treated neurons and found that it was unable to reverse oAβ42 effects on Ca^2+^ activity (oAβ42+inactive Aβcore, 1.77±1.03 ΔF/F_min_) (**Fig. 1**). Finally, inactive Aβcore treatment had no effect on neuronal activity in sAβ42-treated control neurons (sAβ42+inactive Aβcore, 0.95±0.60 ΔF/F_min_) (**Fig. 1**). We thus demonstrated that co-treatment with the Aβcore peptide following application of oAβ42 significantly attenuated oAβ42-induced Ca^2+^ hyperactivity, possibly due to the inhibition of the interaction between Aβ42 and nAChRs.

**Figure 1.**
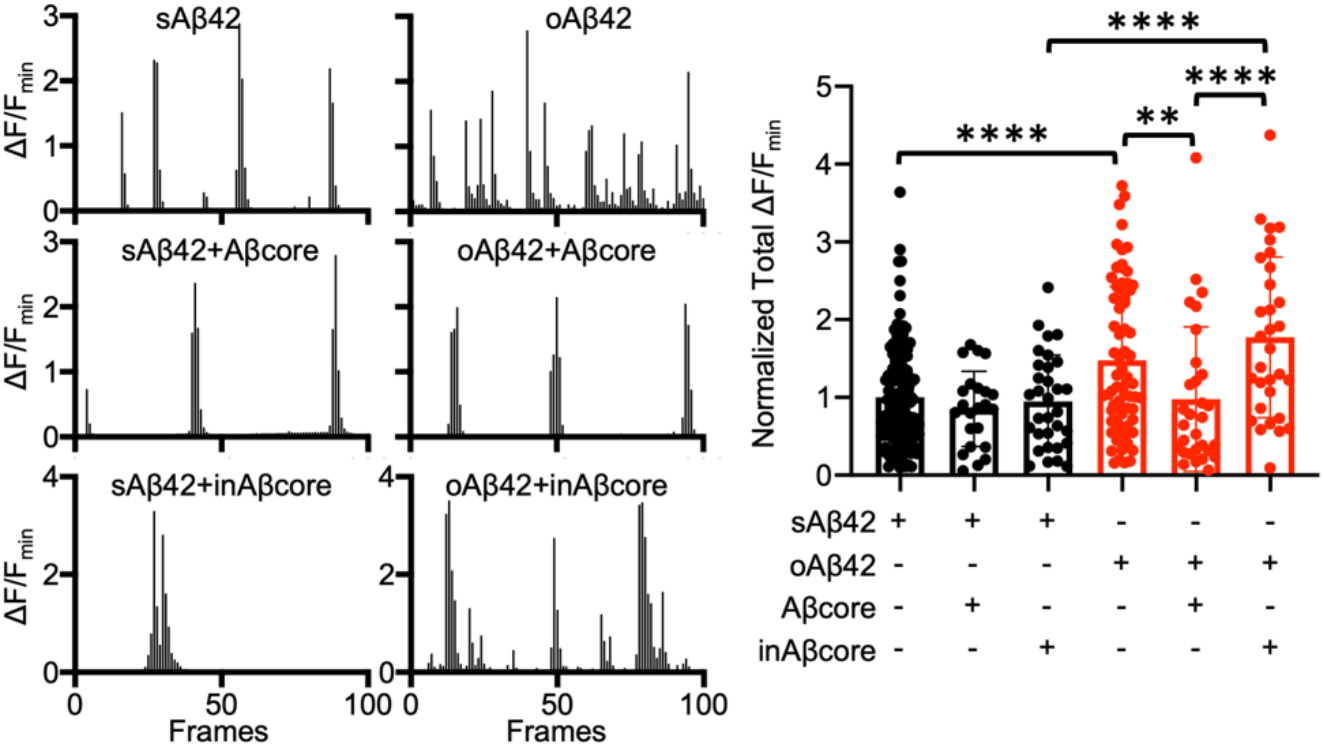
Interaction between Aβ42 and nAChRs is important for Aβ42-induced Ca^2+^ hyperexcitation. Representative traces of GCaMP6f fluorescence intensity in hippocampal cells in each condition and a summary graph of the normalized average of total Ca^2+^ activity in neurons treated with either 250nM sAβ42 (black) or 250nM oAβ42 (red) in the absence or presence of 1μM Aβcore or inactive 1μM Aβcore (inAβcore) (n = number of neurons [sAβ42, n = 127, sAβ42+Aβcore, n = 23, sAβ42+inAβcore, n = 32, oAβ42, n = 71, oAβ42+Aβcore, n = 30, and oAβ42+inAβcore, n = 31], \*\**p* < 0.01 and \*\*\*\**p* < 0.0001 and, one-way ANOVA, Fisher’s LSD Test).

### Aβ selectively interacts with α7 and α4-containing nAChRs, but not α3-containing nAChRs

Many studies support that Aβ can physically interact with α7-, α4- and β2-containing nAChRs in various model systems (17,31,48,49), whereas Aβ is unable to affect α3 and β4-containing receptor function when heterologously expressed in *Xenopus* oocytes (23). Nonetheless, the exact nature of the selective Aβ interaction with nAChRs is not fully defined. To directly measure interactions of Aβ with nAChR subunits, we carried out a series of co-immunoprecipitation (co-IP) analyses in transfected human embryo kidney (HEK293) cells, as described previously (50). Lysates from cells overexpressing human α7-nAChR-GFP receptors were incubated with 2μM Aβ42 for 18 hours and immunoprecipitated with an anti-GFP antibody. We found that the antibody pulled down α7-nAChR-GFP receptors along with Aβ42 (**Fig. 2A**), consistent with previous evidence for an interaction between Aβ42 and α7-nAChR in a neuronal cell line (55). We next expressed mouse α4-nAChR-CFP receptors, and co-IP showed that Aβ42 also interacted with the α4 subunit (**Fig. 2B**). Similar analysis using cells overexpressing human α3-nAChR-GFP receptors yielded no Aβ42 co-immunoprecipitated (**Fig. 2C**). These data suggest Aβ42 can associate with the α4 and α7 subunits but is unable to interact with the α3 subunit.

**Figure 2.**
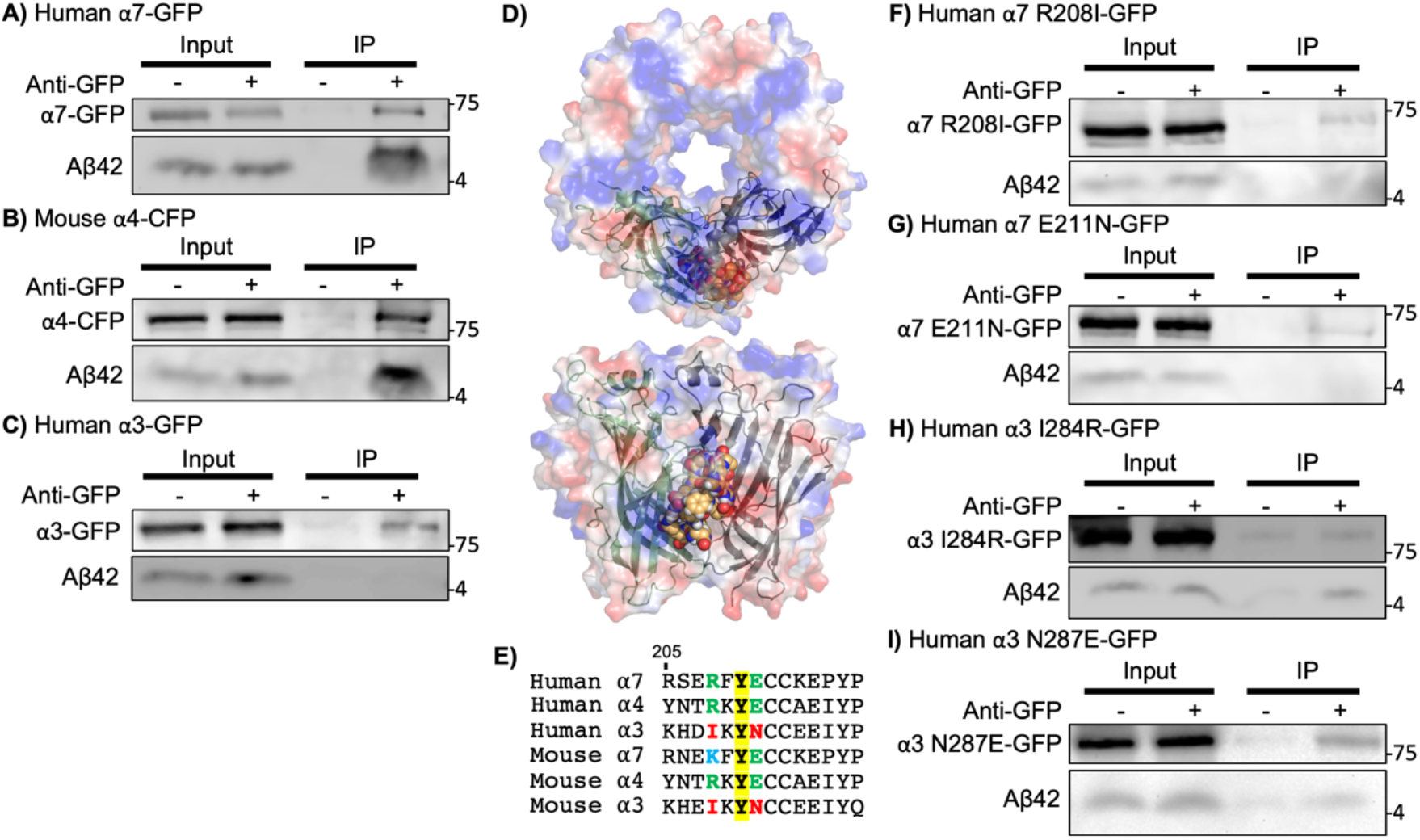
Aβ42 selectively interacts with α7 and α4-containing nAChRs, but not α3-containing nAChRs. **A)** co-IP shows Aβ42 interacts with α7 nAChRs. **B)** co-IP shows Aβ42 binds to α4 nAChRs. **C)** co-IP shows Aβ42 is unable to interact with α3 subunits. **D)** A docking model between N-terminus of Aβ (1-15 amino acids) and pentameric extracellular domains in acetylcholine-binding protein (AChBP) by using CABS-dock server for flexible protein-peptide docking suggests Aβ is likely to sit between two protomers among pentamers. N-terminus of Aβ is shown as a sphere model within vacuum electrostatic model of extracellular domains in AChBP and only two protomers among pentamers are shown as cartoon drawing in top (above) and side views (bottom). **E)** Sequence and numbering of human α7 nAChRs in the loop C region and its alignment with related human and mouse nAChR sequences. Y210 (Bold) is the ligand-binding residue and conserved in all human and mouse α subunits. R208 (Blue) and E211 (Green) are predicted to be critical for interaction with N-terminus of Aβ, which are conserved only in both human and mouse α4 and α7 subunits except mouse α7 receptors that have positive charged Lysine (Light blue), which is similar to positive charged Arginine. However, both mouse and human α3 receptors have uncharged residues in both positions (Red). **F)** Co-IP shows Aβ42 is unable to interact with α7 R208I mutant. **G)** Co-IP shows Aβ42 is unable to bind to α7 E211N mutant. **H)** Co-IP shows Aβ42 can interact with α3 I284R mutant. **I)** Co-IP shows Aβ42 binds to α3 N287E mutant.

Computer simulated docking studies using the homology model of the human α7-nAChRs derived from the X-ray structure of the acetylcholine-binding protein (AChBP) and human Aβ show that the N-terminus of Aβ is predicted to bind to the loop C of the α7 subunit, which is located within the binding interface of two α7 subunits (43,51–54). We used the CABS-dock server for flexible protein-peptide docking (55) to analyze interactions of the α7 nAChR-AChBP chimera (PDB code: 1UW6) (53) and human N-terminus of Aβ (**Fig. 2D**). Three amino acids in the loop C of α7 nAChRs, Arginine 208 (R208), Tyrosine (Y210), and Glutamate 211 (E211), were predicted to be critical for interactions of the α7 subunit with Aβ (**Fig. 2E**). Among them, Y210 is the ligand-binding residue and conserved in all human and mouse α3, α4 and α7 subunits (53) (**Fig. 2E**). Mutation studies have shown that Y210 is essential for acetylcholine binding and Aβ interactions (53,54). Interestingly, both R208 and E211, non-contact residues, are only conserved in human α4 and α7 subunits but not in the α3 subunit (**Fig. 2E**). Importantly, R208 in the human α7 subunit contains positively charged side chain, and mouse α7 subunits contain Lysine (K), a positively charged amino acid, instead of Arginine (**Fig. 2E**). However, both mouse and human α3 subunits contain hydrophobic Isoleucine (I) instead of positively charged Arginine in the human α7 subunit or Lysine in the mouse α7 subunit (**Fig. 2E**). Moreover, E211 in the human α7 subunit contains a negatively charged side chain, which is conserved in both human and mouse α4 and α7 subunits, while both mouse and human α3 subunits include uncharged Asparagine (N) (**Fig. 2E**). Importantly, mutations in R208 and E211 in α7-nAChRs alter the binding affinity of the receptor to acetylcholine (53). Thus, it is possible that these two charged residues are responsible for the nAChR subtype selectivity of Aβ interactions. To test this idea, we generated mutant α7 subunits by substituting R208 for Isoleucine (R208I) or E211 for Asparagine (E211N) to mimic the α3 subunit (**Figs. 2F and 2G**). Co-IP experiments were performed with lysates from HEK293 cells overexpressing human α7-nAChR-R208I-GFP or α7-nAChR-E211N-GFP receptors. We found that Aβ42 was unable to interact with both mutant α7-nAChRs, a loss-of-function effect (**Figs. 2F and 2G**). Furthermore, we made mutant α3 subunits by substituting I284 for Arginine (I284R) or N287 for Glutamate (N287E) to mimic the α7 subunit to test whether these mutants would show gain-of-function effects on the interaction between Aβ42 and the receptors. We carried out Co-IP experiments with lysates from HEK293 cells overexpressing human α3-nAChR-I284R-GFP or α3-nAChR-N287E-GFP receptors. Both mutant receptors were able to pull down Aβ42 (**Fig. 2H and 2I**). These data suggest that the charged Arginine and/or Glutamate residues in the loop C in the α4 and α7 subunits play important roles in the interaction between Aβ and nAChRs, providing direct molecular evidence of how Aβ selectively interacts with α7 and α4-containing receptors, but not α3-containing nAChRs.

### Selective co-activation of α7- and α4β2-nAChRs reverses Aβ-induced reduction of AMPAR surface expression

Aβ has been reported to affect the function of the glutamatergic a-amino-3-hydroxy-5-methyl-4-isoxazolepropionic acid receptors (AMPARs), which are important in synaptic plasticity (56). Several studies suggest Aβ-induced Ca^2+^ hyperexcitation promotes AMPAR endocytosis, which ultimately decreases the surface expression of AMPA receptor subunits GluA1 and GluA2, a cellular mechanism underlying Aβ-induced depression of AMPAR-mediated synaptic transmission (32,57–60). Given that selective co-activation of α7- and α4β2-nAChRs reverses Ca^2+^ hyperexcitation in cultured neurons (32), we examined whether selective co-activation of α7- and α4β2-nAChRs reversed the Aβ effects on surface expression of AMPARs. We measured surface expression of AMPARs by biotinylation after 1μM soluble Aβ42 oligomers (oAβ42) were applied to cultured hippocampal neurons for one hour. Scrambled Aβ42 (sAβ42) was treated in neurons as the control. Consistent with the previous findings (32), oAβ42 treatment reduced surface expression of AMPAR subunits GluA1 and GluA2 (**Fig. 3A-C and Table 1-3**). Next, subtype-specific nAChR agonists, 1μM PNU-282987 (α7), 2μM RJR-2403 Oxalate (α4β2), or 1μM NS-3861 (α3β4) was incubated with oAβ42 or sAβ42 for one hour to activate each nAChR subtype. Activation of either α7- or α4β2- or α3β4-nAChRs singularly was unable to reverse the Aβ effects on GluA1 and GluA2 surface levels (**Fig. 3A and Table 1**). Stimulation of each receptor by themselves also had no effect on GluA1 and GluA2 surface expression in control neurons (**Fig. 3A and Table 1**). Importantly, when we concurrently activated α7- and α4β2-nAChRs for one hour using 1μM PNU-282987 and 2μM RJR-2403 Oxalate, GluA1 and GluA2 surface levels were restored to normal levels in cells treated with oAβ42 (**Fig. 3B and Table 2**). However, stimulation of α7- and α3β4-nAChRs or α4β2- and α3β4-nAChRs was unable to reverse the Aβ effects (**Fig. 3B and Table 2**). In addition, co-stimulation of two nAChR subtypes had no effect on GluA1 and GluA2 surface levels in sAβ42-treated neurons (**Fig. 3B and Table 2**). Next, we activated all three types of nAChRs together by treating neurons with three agonists for one hour and found no neuroprotective effect on Aβ-induced reduction of AMPAR surface levels (**Fig. 3C and Table 3**). Stimulation of α7-, α3β4- and α4β2-nAChRs together was unable to alter surface GluA1 and GluA2 levels in sAβ42-treated control cells (**Fig. 3C and Table 3**). Interestingly, 1μM carbachol, a cholinergic agonist, was also unable to reverse the Aβ effects on AMPAR surface expression (**Fig. 3C and Table 3**). Furthermore, carbachol was sufficient to reduce surface GluA1 but not GluA2 expression in control cells, suggesting global stimulation of acetylcholine receptors may exacerbate the Aβ effects in neurons (**Fig. 3C and Table 3**). This suggests selective co-activation of α7- and α4β2-nAChRs is required to abolish the Aβ effects on AMPAR surface expression.

**Figure 3.**
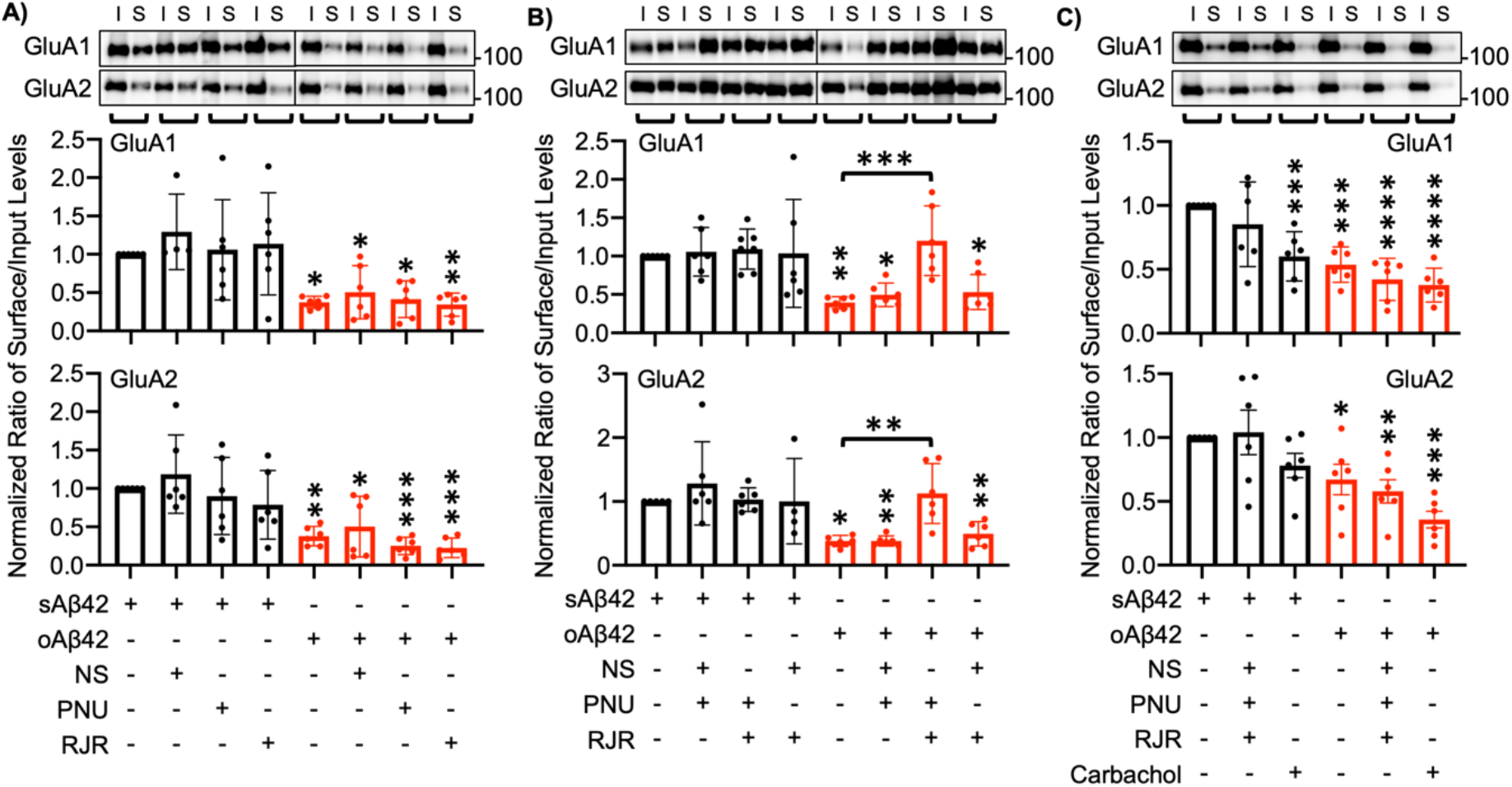
Selective co-activation of α7- and α4β2-nAChRs reverses Aβ-induced reduction of AMPAR surface expression. Representative immunoblots of input (I) and surface (S) levels and quantitative analysis in **A)** Single activation of each nAChRs (n = 6 immunoblots from 3 independent cultures duplicated, \**p* < 0.05, \*\**p* < 0.01, and \*\*\**p* < 0.001, one-way ANOVA, Fisher’s LSD test). **B)** Double activation of each nAChRs (n = 6 immunoblots from 3 independent cultures duplicated, \**p* < 0.05, \*\**p* < 0.01, and \*\*\**p* < 0.001, one-way ANOVA, Fisher’s LSD test). **C)** Triple activation and cholinergic stimulation (n = 6 immunoblots from 3 independent cultures duplicated, \**p* < 0.05, \*\**p* < 0.01, \*\*\**p* < 0.001, and \*\*\*\**p* < 0.0001, one-way ANOVA, Fisher’s LSD test).

**Table 1.** Effects of single agonist application on Aβ-induced reduction of AMPAR surface expression.

| GluA1 | Normalized Ratio of Surface/Input Levels |
| --- | --- |
| sAβ42 | 1.00 |
| sAβ42+NS-3861 | 1.29±0.49 |
| sAβ42+PNU-282987 | 1.06±0.65 |
| sAβ42+RJR-2403 Oxalate | 1.14±0.67 |
| oAβ42 | 0.37±0.08 |
| oAβ42+NS-3861 | 0.51±0.35 |
| oAβ42+PNU-282987 | 0.41±0.24 |
| oAβ42+RJR-2403 Oxalate | 0.34±0.15 |
| GluA2 | Normalized Ratio of Surface/Input Levels |
| sAβ42 | 1.00 |
| sAβ42+NS-3861 | 1.19±0.51 |
| sAβ42+PNU-282987 | 0.90±0.50 |
| sAβ42+RJR-2403 Oxalate | 0.79±0.48 |
| oAβ42 | 0.38±0.13 |
| oAβ42+NS-3861 | 0.50±0.40 |
| oAβ42+PNU-282987 | 0.25±0.11 |
| oAβ42+RJR-2403 Oxalate | 0.23±0.13 |

**Table 2.** Effects of double agonist application on Aβ-induced reduction of AMPAR surface expression.

| GluA1 | Normalized Ratio of Surface/Input Levels |
| --- | --- |
| sAβ42 | 1.00 |
| sAβ42+NS-3861+PNU-282987 | 1.06±0.32 |
| sAβ42+PNU-282987+RJR-2403 Oxalate | 1.09±0.26 |
| sAβ42+RJR-2403 Oxalate+NS-3861 | 1.04±0.70 |
| oAβ42 | 0.39±0.08 |
| oAβ42+NS-3861+PNU-282987 | 0.50±0.15 |
| oAβ42+PNU-282987+RJR-2403 Oxalate | 1.21±0.45 |
| oAβ42+RJR-2403 Oxalate+NS-3861 | 0.53±0.23 |
| GluA2 | Normalized Ratio of Surface/Input Levels |
| sAβ42 | 1.00 |
| sAβ42+NS-3861+PNU-282987 | 1.28±0.65 |
| sAβ42+PNU-282987+RJR-2403 Oxalate | 1.02±0.18 |
| sAβ42+RJR-2403 Oxalate+NS-3861 | 1.00±0.67 |
| oAβ42 | 0.38±0.09 |
| oAβ42+NS-3861+PNU-282987 | 0.38±0.08 |
| oAβ42+PNU-282987+RJR-2403 Oxalate | 1.12±0.47 |
| oAβ42+RJR-2403 Oxalate+NS-3861 | 0.49±0.19 |

**Table 3.** Effects of triple agonist application and cholinergic agonist treatment on Aβ-induced reduction of AMPAR surface expression.

| GluA1 | Normalized Ratio of Surface/Input Levels |
| --- | --- |
| sAβ42 | 1.00 |
| sAβ42+NS-3861+PNU-282987+RJR-2403 Oxalate | 0.85±0.33 |
| sAβ42+Carbachol | 0.6±0.19 |
| oAβ42 | 0.54±0.14 |
| oAβ42+NS-3861+PNU-282987+RJR-2403 Oxalate | 0.42±0.17 |
| oAβ42+Carbachol | 0.38±0.13 |
| GluA2 | Normalized Ratio of Surface/Input Levels |
| sAβ42 | 1.00 |
| sAβ42+NS-3861+PNU-282987+RJR-2403 Oxalate | 1.04±0.43 |
| sAβ42+Carbachol | 0.78±0.23 |
| oAβ42 | 0.67±0.29 |
| oAβ42+NS-3861+PNU-282987+RJR-2403 Oxalate | 0.58±0.22 |
| oAβ42+Carbachol | 0.36±0.16 |

### Co-activation of α7- and α4β2-nAChRs reverses Aβ-induced impaired AMPAR phosphorylation and synaptic plasticity

Several studies suggest Aβ-induced Ca^2+^ hyperexcitation elevates activity of Ca^2+^-dependent phosphatase, calcineurin, which, in turn, will promote AMPAR endocytosis via dephosphorylation of AMPAR subunit GluA1 at serine 845, a residue that plays a crucial role in AMPAR surface expression during synaptic plasticity (32,57,58). In fact, previous studies reveal that Aβ reduces AMPAR GluA1 phosphorylation at serine 845 (pGluA1), which is strongly associated with disrupted LTP in AD (32,57,60). Consistently, hippocampal LTP can be blocked by either direct exogenous Aβ application at high levels or abnormally high levels of Aβ produced from AD transgenic mouse models (57,58,61–64). This may contribute to AD-associated synaptic dysfunction and memory deficits (6). Given that co-activation of α7- and α4β2-nAChRs was sufficient to restore normal AMPAR surface levels (**Fig. 3B**) and neuronal Ca^2+^ activity in Aβ-treated cultured neurons (32), we examined whether co-stimulation of these receptors reversed the Aβ effects on AMPAR phosphorylation and LTP. First, we treated cultured hippocampal neurons with 1μM oAβ42 or sAβ42 for one hour and measured basal pGluA1 levels (**Fig. 4A**). As shown before (32), oAβ42 treatment decreased pGluA1 compared with the sAβ42-treated control (sAβ42, 1.00 and oAβ42, 0.48±0.25, *p* = 0.0424) (**Fig. 4A and 4B**). Significantly, Aβ-induced reduction of pGluA1 was reversed by co-stimulation of α7- and α4β2-nAChRs when we treated neurons with 1μM PNU-282987 and 2μM RJR-2403 Oxalate for one hour (oAβ42+agonists, 1.27±0.91, *p* = 0.003) (**Fig. 4A and 4B**). However, co-activation of these receptors in control cells had no effect on pGluA1 levels (sAβ42+agonists, 0.92±0.36) (**Fig. 4A and 4B**). We next treated neurons with a glycine-based media, well-established to induce a form of chemical LTP (cLTP), as shown previously (65). We treated neurons with 1μM oAβ42 or sAβ42 for one hour, induced cLTP, and measured pGluA1 levels (**Fig. 4A**). As shown previously (65), following cLTP induction, pGluA1 levels were significantly elevated in control neurons, an indication of LTP expression (**Fig. 4A**). However, pGluA1 levels were significantly lower in oAβ42-treated neurons compared with sAβ42-treated control cells after cLTP induction (sAβ42, 1.00 and oAβ42, 0.55±0.33, *p* = 0.0075) (**Figs. 4A and 4C**). Importantly, normal pGluA1 levels were restored when we activated both α7- and α4β2-nAChRs and induced cLTP in oAβ42-treated neurons (oAβ42+agonists, 0.91±0.27, *p* = 0.0359) (**Fig. 4A and 4C**). However, co-activation of these receptors in control cells had no effect on pGluA1 levels following cLTP induction (sAβ42+agonists, 1.14±0.61) (**Fig. 4A and 4C**). Thus, co-activation of α7- and α4β2-nAChRs was sufficient to reverse the Aβ effects on AMPAR phosphorylation and cLTP.

**Figure 4.**
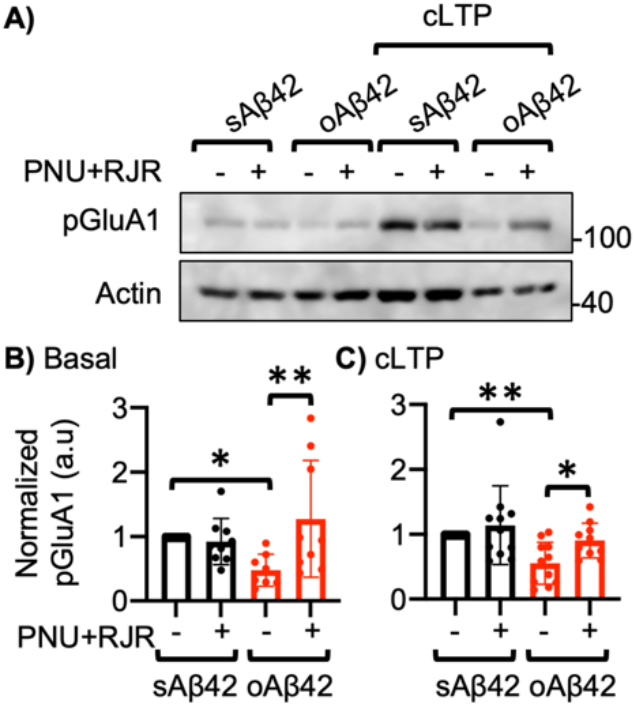
Co-activation of α7- and α4β2-nAChRs reverses Aβ-induced impaired AMPAR phosphorylation and synaptic plasticity. **A)** Representative immunoblots of pGluA1 levels in each condition. **B)** Quantitative analysis of pGluA1 levels under the basal condition in each condition (n = 9 immunoblots from 4 independent cultures, \**p* < 0.05 and \*\**p* < 0.01, one-way ANOVA, Fisher’s LSD test). **C)** Quantitative analysis of pGluA1 levels following cLTP induction in each condition (n = 11 immunoblots from 5 independent cultures, \**p* < 0.05 and \*\**p* < 0.01, one-way ANOVA, Fisher’s LSD test).

## Discussion

Current therapeutic approaches to AD suffer from lack of specificity and poor efficacy. Preclinical approaches based on altering Aβ have failed in clinical trials. Consequently, novel approaches are being explored, including targeting receptors regulated by Aβ. Nicotinic receptors have emerged as potential targets for reversing cognitive deficits in AD (66), owing to their noted potent regulation by Aβ, but there remains a need to determine the roles of specific nAChR subtypes in AD. In this study, we demonstrate that the interaction between Aβ and nAChRs plays an important role in Aβ-induced alteration of synaptic and neuronal activity. We provided further evidence for Aβ’s selective interaction with α7- and α4-containing nAChRs but not α3-containing receptors, and selective stimulation of α7- and α4-containing nAChRs was shown to be neuroprotective against the Aβ effects on synaptic function. Notably, we identified two key amino acids, Arginine and Glutamate, present in the loop C of the α7 and α4 subunits, but not the α3 subunit, that are important for interaction with Aβ, providing a molecular mechanism for Aβ’s selective inhibition of α7- and α4β2-nAChRs.

Based on the present findings, we propose the following model (**Fig. 5**). Given that nAChRs are more prominently expressed in inhibitory interneurons in the hippocampus, soluble Aβ42 oligomers selectively interact with α7- and α4β2-nAChRs but not α3β4-nAChRs and reduce neuronal activity in inhibitory cells, leading to a decrease in the release of GABA onto hippocampal excitatory neurons (**Fig. 5A**). This is supported by our previous work in which stimulation of GAB_A_A receptors is sufficient to reverse Aβ42-induced Ca^2+^ hyperactivity in cultured neurons (32). Excitatory cells will thus have increased neuronal Ca^2+^ activity, consequently elevating the activity of calcineurin (32) (**Fig. 5A**) and other Ca^2+^ signaling pathways. This promotes the dephosphorylation of the AMPAR subunit, GluA1, which allows for AMPAR endocytosis, resulting in an overall decrease of AMPAR surface expression (**Fig. 5A**). This ultimately contributes to disruption of LTP (**Fig. 5A**) and may lead to cognitive decline. As Aβ42 inhibits both α7- and α4β2-nAChRs but not α3β4-nAChRs, co-stimulation of α7- and α4β2-nAChRs by using selective agonists can reverse the Aβ effects on synapses by restoring normal activity of both hippocampal inhibitory and excitatory cells (**Fig. 5B**). With restoration of normal Ca^2+^ activity, calcineurin activity decreases, leading to AMPAR dephosphorylation and decreased AMPAR endocytosis, ultimately restoring normal LTP (**Fig. 5B**). Given that Aβ42 inhibits both α7- and α4β2-nAChRs, stimulation of each receptor by themselves has no neuroprotective effect (**Fig. 3A**).

**Figure 5.**
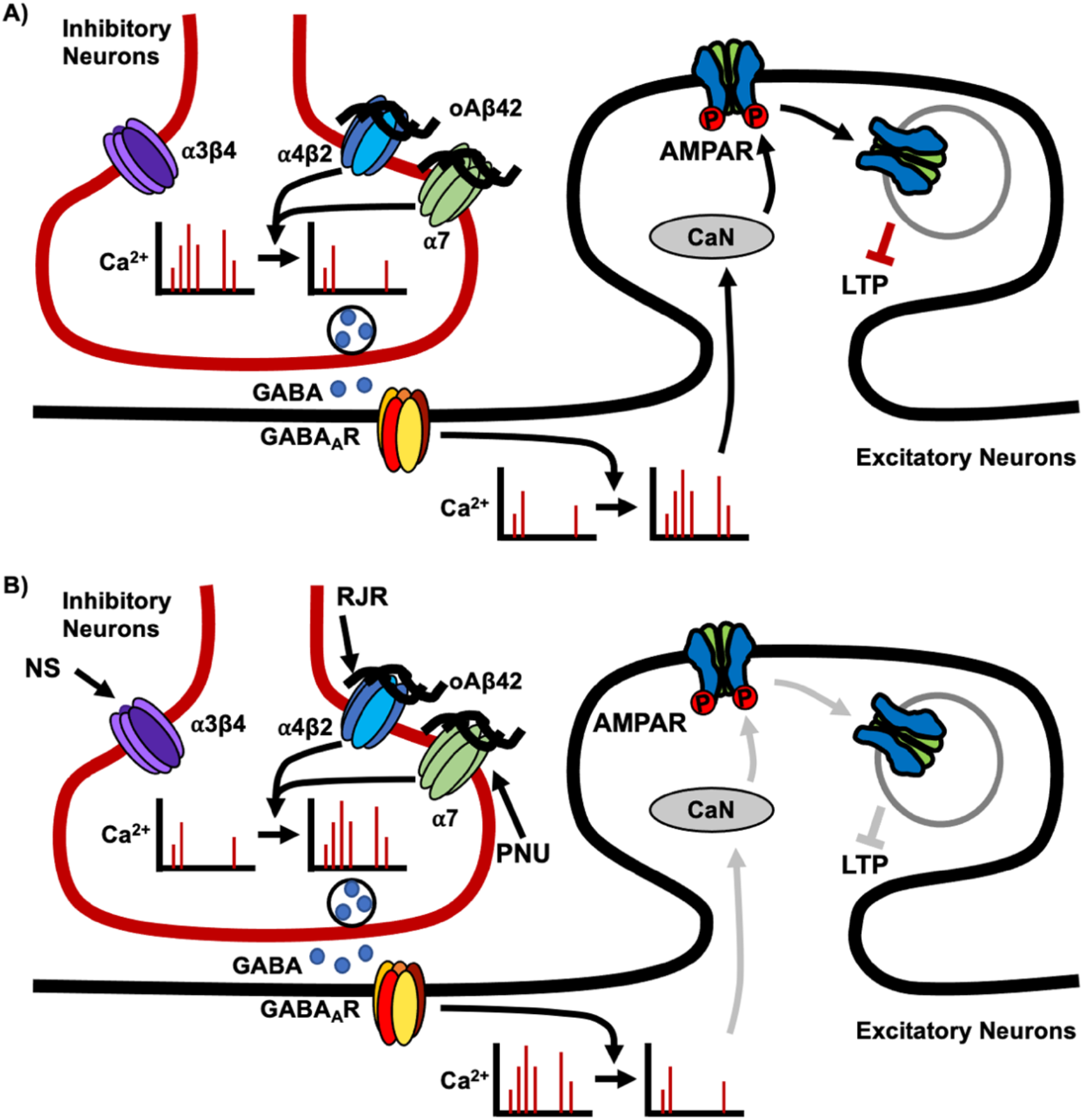
Schematic model. **A**) **Impact of Aβ oligomers.** In the hippocampus, nAChRs are prominently expressed on inhibitory interneurons, thus, selective binding of soluble Aβ42 oligomers (oAβ42) to α7- and α4β2-nAChRs but not α3β4-nAChRs, reduces neuronal activity in inhibitory cells, leading to a decrease in the release of GABA onto hippocampal excitatory neurons. Consequently, excitatory cells have increased frequency of Ca^2+^ transients, resulting in elevated calcineurin (CaN) activity. CaN dephosphorylates the AMPAR subunit, GluA1, promoting AMPAR endocytosis and resulting in an overall decrease of AMPAR surface expression. This ultimately contributes to disruptions of LTP. **B) Reversal of Aβ-induced synaptic and neuronal dysfunction by co-stimulation with α7- and α4β2-nAChRs agonists.** As Aβ42 inhibits both α7- and α4β2-nAChRs but not α3β4-nAChRs, co-stimulation of α7- and α4β2-nAChRs by selective agonists, PNU-282987 (PNU) and RJR-2403 Oxalate (RJR), can restore normal activity of both hippocampal inhibitory and excitatory cells, reversing Aβ-induced synaptic dysfunction. This restoration of normal Ca^2+^ activity prompts a decrease in calcineurin activity, leading to a decrease in AMPAR dephosphorylation and AMPAR endocytosis, ultimately restoring normal LTP. However, an agonist for α3β4-nAChRs, NS-3861 (NS), does not appear to have neuroprotective effects. Moreover, non-specific stimulation of nAChRs by using three agonists together or carbachol is unable to reverse the Aβ effects on neuronal activity and synaptic function, emphasizing the importance of selective co-stimulation of nAChRs as potential therapeutic approaches.

Our co-IP experiments using mutant α7- and α3-containing receptors confirm that the charged Arginine and Glutamate residues in the loop C of α7- and α4-containing nAChRs are critical for the interaction of Aβ with these receptors. In fact, a previous study identifies that R208 and E211 in the loop C region of the α7-nAChRs interact with juxtapose residues with the same charge, in which R208 pairs with Lysine 167 and E211 pairs with Glutamate 184 and Aspartic acid 186 (53). These interactions create electrostatic repulsion that likely regulate binding affinity of agonists to the receptor (53). Interestingly, a loss of one of these charged residues in the α7 subunit (R208I or E211N) disrupts the interaction of mutant α7-containing receptors with Aβ. Where α7-nAChRs are primarily present as homopentamers (53), the electrostatic repulsion is established within the α7 subunits, and thus mutations of R208 or E211 disrupt this electrostatic repulsion, which may contribute to loss of binding to Aβ. Unlike the α7 subunits, a gain of one of these charged residues in the α3 subunit (I284R or N287E) enabled the receptors to associate with Aβ. In contrast to homomeric α7-nAChRs, α4- and α3-containing receptors exist as heteropentamers with β subunits (67,68). Interestingly, it seems that Glutamate in the loop C of the α4 subunit may pair with a negatively charged residue in the β subunits, providing the electrostatic repulsion similar to α7 homopentamers (67). Thus, the β subunits may contribute to the binding affinity of the receptor to Aβ in α4β2-, α3β4-, and α7β2-nAChRs, another Aβ-interacting receptors expressed in hippocampal inhibitory neurons (69). Given that the α3 subunit contains non-charged amino acids in the loop C, a gain of charged residues in the loop C of α3-containing receptors may increase affinity of the receptors for Aβ42. In addition, other computation modeling studies show E211 in the loop C of the α7 subunit is able to interact with Aβ42 and further suggest that the similar interactions apply to α4β2-AChRs (70,71). In sum, these findings suggest the charged residues, Arginine and Glutamate, in the loop C of the α7 and α4 subunits are critical for Aβ interaction and its effect on the receptors.

The current study focuses on the differential effects of Aβ on nAChR subtypes that lead to alterations in neuronal excitability and synaptic function. Consistent with previous findings using acetylcholinesterase inhibitors (74), we found that non-selective cholinergic activation exacerbates Aβ effects on hippocampal neurons, pointing to confounding effects of activating α3β4-nAChRs, which may result in unexpected side effects, including imbalance of excitation and inhibition in hippocampal circuits (72). Therefore, acetylcholinesterase inhibitors are not very effective in slowing AD progression due to non-selective stimulation of acetylcholine receptors (73,74). There are also discrepancies involving the use of nicotine treatment to stimulate nAChRs to alter cognitive function. For example, nicotinic agonists have been found to improve performance in a variety of memory tasks in rodents and non-human primate studies (66), while several other studies have failed to find significant enhancement of learning and memory by nicotine treatment (75). Part of the discrepancy may lay in the antagonistic effect of prolonged nicotine exposure, owing to induction of nAChR inactivation (desensitization). The extent to which nAChR subtype-selective agonists drive receptor inactivation, and their impact on intracellular signaling in Aβ neurotoxicity remains to be determined (76). Nonetheless, nAChR agonists have consistently suggested promising approaches in the treatment of AD (77). However, clinical trials thus far have been challenged by adverse effects or minimal improvement (78). In particular, stimulating only one type of nAChRs by using a subtype-specific agonist was found to either enhance cognitive performance or have no beneficial effect. For instance, selective α7-nAChR agonists have been reported to improve cognition in a variety of animal models (79–81), while another study has found they have almost no beneficial effect on learning and memory in mice (82). An α4β2-nAChR agonist alone can improve working memory only in young rats but not older animals (83). It is thus not yet clear whether single activation of specific nAChR subtypes provides optimal efficacy in AD (77). We have shown that activation of either α7- or α4β2-nAChRs singularly had no effect on Aβ-induced hyperexcitation (32) or α7- and α4β2-nAChR Aβ-induced reduction of AMPAR surface expression (**Fig. 3A**). By contrast, selective co-activation of α7- and α4β2-nAChRs was sufficient to reverse Aβ-induced neuronal hyperexcitation (32) and synaptic dysfunction (**Fig. 3B and 4**). Several subtype-specific agonists have been developed for clinical trials, but co-activation of nAChRs has not been applied for clinical trials yet (77), thus the current study may lead to an innovative and novel therapeutic strategy for AD.

## Experimental procedures

### Mouse hippocampal neuron culture

Mouse hippocampal neuron cultures were prepared as described previously (32,47,84–86). Hippocampi were isolated from postnatal day 0 (P0) CD-1 (Charles River) mouse brain tissues and digested with 10U/mL papain (Worthington Biochemical Corp. Lakewood, NJ). For Ca^2+^ imaging, mouse hippocampal neurons were plated on poly-D-lysine-coated glass bottom dishes (5 × 10^5^ cells) and imaged on day *in vitro* (DIV) 12-14. For biotinylation assays and cLTP, hippocampal neurons were plated in 6 cm dishes (3 × 10^6^ cells) and used on DIV 14. Cells were grown in Neurobasal medium (Life Technologies, Carlsbad, CA) with B27 supplement (Life Technologies, Carlsbad, CA), 0.5mM Glutamax (Life Technologies) and 1% penicillin/streptomycin (Life Technologies). Colorado State University’s Institutional Animal Care and Use Committee reviewed and approved the animal care and protocol (978).

### Reagents

Soluble Aβ42 oligomers or soluble scrambled Aβ42 oligomers were prepared as previously described (32). One mg of lyophilized human Aβ42 (Anaspec) or scrambled Aβ42 (Anaspec) was dissolved in 1mL of 1,1,1,3,3,3-hexafluoro-2-propanol (HFIP) (Fisher Scientific) to prevent aggregation, portioned into 10μg aliquots, air-dried and stored at −80°C. For use in experiments, an aliquot was thawed at room temperature, and then dissolved in 100% DMSO then diluted into PBS to make a 100μM solution. The solution was incubated for 16 hours at 4°C and then diluted to a final concentration for use in experiments. The following agonists were used in this study: 1μM PNU-120596 (Alomone labs), 2μM RJR-2403 Oxalate (Alomone labs), 1μM NS-3861 (Tocris Bioscience), and 1μM Carbamoylcholine chloride (carbachol) (Tocris Bioscience). Aβcore (YEVHHQ) and inactive Aβcore (SEVAAQ) peptides were prepared as described previously (43).

### HEK293 Cell culture and transfection

HEK293 cells were grown in DMEM medium with fetal bovine serum (Life Technologies) and 1% penicillin/streptomycin (Life Technologies) and transfected by jetPRIME® DNA and siRNA transfection reagent (Polyplus) according to the manufacturer’s protocol. Cells used for each experiment were from more than three independently prepared cultures.

### DNA plasmids and mutagenesis

Human pcDNA3.1-CHRNA7-mGFP was a gift from Henry Lester (Addgene plasmid # 62629; http://n2t.net/addgene:62629; RRID:Addgene_62629). Mouse nAChR alpha4 CFP was a gift from Henry Lester (Addgene plasmid # 15244; http://n2t.net/addgene:15244; RRID:Addgene_15244). Human α3-nAChR-GFP was obtained from Sino Biological (HG29719-ACG). Human α7-nAChR R208I-GFP and E211N-GFP were generated from pcDNA3.1-CHRNA7-mGFP and human α3-nAChR I284R-GFP and N287E-GFP were generated from human α3-nAChR-GFP by PCR based QuikChange® Site-Directed Mutagenesis Kit (Agilent) according to the manufacturer’s protocol. The following primers were used for mutagenesis; R208I (5-GGA ATC CCC GGC AAG AGG AGT GAA ***ATA*** TTC TAT GAG TGC TGC-3 and 5-CCT TAG GGG CCG TTC TCC TCA CTT ***TAT*** AAG ATA CTC ACG ACG-3), E211N (5-AGG TTC TAT ***AAC*** TGC TGC AAA GAG CCC TAC CCC GAT GTC-3 and 5-TCC AAG ATA ***TTG*** ACG ACG TTT CTC GGG ATG GGG CTA CAG-3), I208R (5-CCA GGC TAC AAA CAC GAC ***CGC*** AAG TAC AAC TGC TGC −3 and 5-GCA GCA GTT GTA CTT ***GCG*** GTC GTG TTT GTA GCC TGG −3) and N211E (5-GGC TAC AAA CAC GAC ATC AAG TAC ***GAG*** TGC TGC GAG GAG −3 and 5-CTC CTC GCA GCA ***CTC*** GTA CTT GAT GTC GTG TTT GTA GCC −3); Bold and italic nucleotides indicate mutations introduced.

### Co-immunoprecipitation

Co-IP experiments were performed using a Co-IP kit (Pierce) following the manufacturer’s protocol with samples from three independently prepared culture as carried out previously (50). Plasmids expressing human α7-nAChR-GFP, human α7-nAChR R208I-GFP, human α7-nAChR E211N-GFP, human α4-nAChR-CFP, human α3-nAChR-GFP, α3-nAChR I284R-GFP or α3-nAChR N287E-GFP were transfected in HEK293 cells. 2μM soluble Aβ42 were added to total cell lysates (200μl), and 20μl of cell lysates with Aβ42 were collected as the input. 180μl of cell lysates incubated with Aβ42 were pulled down with anti-GFP antibody. Immunoprecipitated samples were applied to immunoblots.

### Computational modeling

Computational peptide docking of N-terminus of Aβ into the α7-nAChR-AChBP (the acetylcholine-binding protein) chimera (53) was performed using the CABS-dock server for flexible protein-peptide docking (55). For the α7-nAChR-AChBP chimera, the X-ray crystallographic structure encompassing two adjacent α subunits containing the ligand binding domain, equivalent in all five sites in the pentameric receptor, was used in a flexible protein-peptide docking.

### GCaMP Ca^2+^ Imaging

GCaMP Ca^2+^ imaging was carried out by the previously reported method (32,47). DIV 4 neurons were transfected with pGP-CMV-GCaMP6f (a gift from Douglas Kim, Addgene plasmid # 40755; http://n2t.net/addgene:40755 ; RRID:Addgene_40755) (87) for imaging hippocampal pyramidal cells or pAAV-mDlx-GCaMP6f-Fishell-2 (a gift from Gordon Fishell, Addgene plasmid # 83899; http://n2t.net/addgene:83899; RRID:Addgene_83899) (88) for imaging interneurons by using Lipofectamine 2000 (Life Technologies) according to the manufacturer’s protocol. Neurons were imaged DIV 12-14. The transfection efficiency was around 2% and no obvious cellular toxicity has been observed. Neurons were grown in Neurobasal Medium without phenol red (Life Technologies) and with B27 supplement (Life Technologies), 0.5mM Glutamax (Life Technologies) and 1% penicillin/streptomycin (Life Technologies) for 8-10 days after transfection and during the imaging. Glass-bottom dishes were mounted on a temperature-controlled stage on an Olympus IX73 microscope and maintained at 37°C and 5% CO_2_ using a Tokai-Hit heating stage and digital temperature and humidity controller. For GCaMP6f, the images were captured with a 10 ms exposure time and a total of 100 images were obtained with a 500 ms interval. F_min_ was determined as the minimum fluorescence value during the imaging. Total Ca^2+^ activity was obtained by 100 values of ΔF/F_min_ = (F_t_ – F_min_)/F_min_ in each image, and values of ΔF/F_min_ < 0.1 were rejected due to bleaching. Ten to twenty neurons were used for imaging in each individual experiment, and one individual neuron was assayed in an image.

### Surface Biotinylation

Surface biotinylation was performed according to the previous studies (32,47,84–86). Cells were washed with ice-cold PBS containing 1mM CaCl_2_ and 0.5mM MgCl_2_ and incubated with 1mg/ml Sulfo-NHS-SS-biotin (Thermo Scientific) for 15 min on ice. Following biotin incubation, neurons were washed with 20 mM glycine to remove the excess of biotin, and cells were lysed in 300μl RIPA buffer for one hour. 10% of total protein was separated as an input samples, and protein lysates were incubated overnight with streptavidin-coated beads (Thermo Scientific) at 4°C under constant rocking. The beads containing surface biotinylated proteins were separated by centrifugation. Biotinylated proteins were eluted from streptavidin beads with SDS loading buffer. Surface protein fractions and their corresponding total protein samples were analyzed by immunoblots.

### cLTP

cLTP was performed by a modification of the previously described method (65). Cells were washed with Mg^2+^ free buffer (150 mM NaCl, 5 mM KCl, 1 mM CaCl_2_, 33mM glucose, 10mM HEPES [pH 7.4], 20μM bicuculline, 1μM strychnine), treated with 200μM glycine in Mg^2+^ free buffer for 5 min at 37°C, and returned to Mg^2+^ buffer (Mg^2+^ free buffer and 2mM MgCl_2_) for 20 min at 37°C. Cells were then lysed with 100μl RIPA buffer, and cell lysates was used for immunoblots.

### Immunoblots

Protein samples for biotinylation and cLTP were loaded on 10% glycine-SDS-PAGE gel. Co-IP samples were loaded 16% Tricine-SDS-PAGE gel as described previously (89). SDS-PAGE gels were transferred to nitrocellulose membranes. The membranes were blocked (5% powdered milk) for 1 h at room temperature, followed by overnight incubation with the primary antibodies at 4°C. The primary antibodies consisted of anti-GluA1 (Millipore, 1:2000), anti-GluA2 (Abcam, 1:2000), anti-phosphorylated GluA1 S845 (Millipore, 1:1000), anti-GFP (Torrey Pines, 1:2000), anti-Aβ (6E10, Covance, 1:2000), and anti-actin (Abcam, 1:2000) antibodies. Membranes were subsequently incubated by secondary antibodies for 1 h at room temperature and developed with Enhanced Chemiluminescence (ECL) (Thermo Fisher Scientific, Waltham, MA). Protein bands were quantified using ImageJ (https://imagej.nih.gov/ij/). Immunoblots were at least duplicated for quantitative analysis.

### Statistics

Statistical comparisons were analyzed with the GraphPad Prism 8 software. Unpaired two-tailed Student t-tests were used in single comparisons. For multiple comparisons, one-way ANOVA followed by Fisher’s Least Significant Difference (LSD) test was used to determine statistical significance. Results are represented as mean ± Standard Deviation (SD), and *p* < 0.05 was considered the minimum for statistical difference.

## Acknowledgments

We thank members of the Kim laboratory for their generous support. In particular, we thank Rosaline Danzman, Luis Gomez Wulschner, Thomas McMillan, and Caleb Wipf for initiating the project. We also thank Drs. Michael Tamkun, Jozsef Vigh, Frederic Hoerndli, and Susan Tsunoda for helpful discussion and providing reagents. We specially thank Dr. SeungBeom Hong at King Abdullah University of Science and Technology for his help in structural analysis.

## Data availability

All data are contained within the manuscript.

## Funding and additional information

This work is supported by the University of Hawai’i Foundation Preclinical Alzheimer’s Research Fund (RAN), the pilot program from the Colorado Clinical and Translational Sciences Institute (SK), College Research Council Shared Research Program from Colorado State University (SK), the Boettcher Foundation (SK), The Barry Goldwater Scholarship and Excellence in Education Foundation (JPR), Astronaut Scholarship Foundation (JPR), and Student Experiential Learning Grant from Colorado State University (JPR and SAS).

